# Microcalcifications can either trigger or suppress breast precancer malignancy potential according to the mineral type in a 3D tumor model

**DOI:** 10.1101/2023.02.20.529220

**Authors:** Amit Cohen, Lotem Gotnayer, Dina Aranovich, Netta Vidavsky

## Abstract

Most early breast precancer lesions, termed ductal carcinoma *in situ* (DCIS), contain microcalcifications (MCs), which are calcium-containing pathological minerals. The most common type of MCs is calcium phosphate crystals, mainly carbonated apatite; it is associated with either benign or malignant lesions. *In-vitro* studies indicate that the crystal properties of apatite MCs can affect breast cancer progression. A less common type of MCs is calcium oxalate dihydrate (COD), which is almost always found in benign lesions. We developed a 3D tumor model of multicellular spheroids of human precancer cells containing synthetic MC analogs that link the crystal properties of MCs with the progression of breast precancer to invasive cancer. We show that apatite crystals induce proliferation and Her2 overexpression in DCIS cells. This tumor-triggering effect is increased when the carbonate fraction in the MCs decreases. COD crystals, in contrast, do not induce proliferation and reduce Her2 expression, even compared with control spheroids with no added MC analogs. This finding suggests that COD is not randomly located only in benign lesions—it may actively contribute to suppressing precancer progression in its surroundings. Our model provides an easy-to-manipulate platform to better understand the interactions between breast precancer cells and MCs. A better understanding of the effect of the crystal properties of MCs on precancer progression will potentially provide new directions for better precancer prognosis and treatment.

## 1. Introduction

Most breast cancers develop in the mammary glands as the epithelial cells lining the milk duct become cancerous, fill the duct lumen, and invade the surrounding tissue^1^. In ductal carcinoma *in situ* (DCIS), a common non-palpable early breast precancer tumor, the cancer cells are still confined to the duct lumen. Because it is impossible to predict whether DCIS will eventually become invasive cancer, it is treated as such. When screening for breast cancer via mammography, one of the most common features observed in DCIS cases that improve early detection is microcalcifications (MCs)^2,3^. MCs are calcium-containing pathological minerals whose appearance in mammograms is associated with malignancy and poorer breast cancer prognosis^4,5^. DCIS is the breast cancer subtype most associated with MCs. In DCIS patients, mammographic MCs are associated with increased local cancer reoccurrence^6^ and overexpression of human epidermal growth factor receptor 2 (Her2)^7^.

Breast MCs are heterogeneous in composition and morphology, and several calcium-containing minerals have been found in breast tissues^8,9^. A well-studied mineral often observed in breast tissues is calcium phosphate, mostly in the form of carbonated apatite, found in benign or malignant breast lesions^10–12^. Biological apatite crystals vary in composition and contain various ions in addition to the calcium, phosphate, and hydroxyl ions found in stoichiometric hydroxyapatite. Alterations in the apatite composition correlate with lesion malignancy. For example, low carbonate and high sodium apatites are associated with more invasive lesions^11,13,14^. Calcium oxalate dihydrate (COD) is another type of breast MC that is less common than apatite^13,15^. Unlike apatite MCs, which are associated with either benign or malignant lesions, COD MCs are almost always found in benign lesions and are rarely found in DCIS ^12^,^14^,^16–20^.

Currently, breast cancer screening, diagnosis, and prognosis only utilize the appearance of MCs in mammograms and histopathology to inform about the stage of the disease. However, the correlation between MC crystal properties and malignancy raises the possibility that the chemical properties of MCs can improve breast cancer diagnosis and prognosis^21^. Additionally, several *in-vitro* and *in-vivo* studies have examined how the crystal properties of apatite affect breast cancer progression. *S*tudies of murine and human mammary epithelial cell lines cultured as monolayers indicated that hydroxyapatite enhances mitogenesis and cell migration, whereas calcium oxalate monohydrate (COM) does not^10,22^. Interestingly, 3D polymeric scaffolds containing hydroxyapatite crystals promoted the cancerous behavior of human precancer cells and invasive breast cancer cell lines, compared with mineral-free scaffolds^23–25^. In these systems, hydroxyapatite promoted the invasiveness of human DCIS cells^25^, and the crystallinity and particle size of hydroxyapatite crystals affected the malignancy potential of invasive breast cancer cells^24^. Human invasive cell lines cultured on hydroxyapatite surfaces with changing properties altered their cancer behavior according to the hydroxyapatite carbonate concentration, surface area, and roughness^26^. Even though evidence supports the triggering effect of calcium phosphate on breast cancer progression, to the best of our knowledge, the effect of COD on human breast precancer cell progression has not yet been studied, probably because COD crystals are primarily found in benign lesions and are considered inert regarding disease progression.

*In-vitro* tumor models are a practical platform to study mineral-precancer cell interactions because they allow dynamic process monitoring while mimicking the conditions of breast tumors^27^. Although clinical samples provide a more realistic view of the tumor, they are heterogeneous and complex, making it challenging to address mechanistic questions and determine the effect of specific parameters on disease progression^28^. Precancer cells can be cultured as 3D multicellular spheroids that maintain cell-cell and cell-extracellular matrix interactions while maintaining the cells in the tumor microenvironmental context and mimicking the DCIS tumor *in vitro*^29–31^. Such multicellular spheroids better simulate how cells behave *in vivo* than do 2D cultures^32^.

Here, we developed a 3D-engineered tumor model of breast precancer cells cultured as multicellular spheroids embedded with synthetic MC analogs to mimic the 3D microenvironment of mammary ducts that contain MCs. Specifically, we used precancer cells of the human cell line MCF10DCIS.com, which, *in vivo*, induce DCIS. These cells are derived from MCF10A cells, a normal human epithelial cell line isolated from a patient with fibrocystic breasts^33^ and are estrogen receptor (ER) negative as well as progestogen receptor (PR) and human epidermal growth factor receptor (Her2) positive^34^. MCF10DCIS.com cells were shown to spontaneously deposit MC-like mineral particles after ten days of culture as multicellular spheroids^35^. In the system that we developed, we did not wait for spontaneous mineral deposition but instead, added synthetic minerals with controlled properties to the spheroids. We used mineral types that are physiologically relevant such as apatite with high and low carbonate content associated with benign and malignant lesions, respectively, and COD, which is always found in benign lesions.

## 2. Experimental

### 2.1 Mineral synthesis

All materials were purchased from Sigma Aldrich (St. Louis, MO, USA), except the calcium chloride, which was purchased from Fisher Scientific (Loughborough, UK). High carbonate apatite (HCA) and COD were synthesized using established protocols^36,37^. Briefly, for HCA synthesis, a phosphate solution ((NH_4_)_2_HPO_4_: 10.0 g, (NH_4_)CO_3_: 2.5 g, NH_4_OH: 25 mL, H_2_O:150 mL) and a calcium solution (Ca(NO_3_)_2_·4H_2_O: 23.6 g, NH_4_OH: 50 mL, H_2_O: 150 mL) were prepared. The solutions were mixed and heated to 85°C. The solution was stirred for one hour at 85°C, cooled to room temperature (RT), and centrifuged; then the precipitate was washed with deionized water.

For COD synthesis, a calcium chloride solution (CaCl_2_: 7.10 mg, H_2_O: 32 mL) and a sodium citrate solution (Na_3_C_6_H_5_O_7_: 1176 mg, H_2_O: 400 mL) were stirred at 100 rpm at RT. Then, a sodium oxalate solution (Na_2_C_2_O_4_: 107 mg, H_2_O: 16 mL) was added to the solution, and they were stirred continuously at 100 rpm at RT for 15 min. A precipitate was obtained after centrifugation and washing with deionized water and ethanol.

### 2.2 Mineral characterization

#### Fourier Transform Infrared (FTIR) spectroscopy

All mineral particles were analyzed using FTIR spectroscopy (Nicolet iS5, Thermo Fischer). First, 1 mg from each sample was mixed with 200 mg of KBr until the powder was homogeneous; then a pellet was made with a compressor rated at 7 tons for 5 minutes. The Nicolet iS5 instrument was operated in the spectral range of 400–4000 cm^-1^ with 32 scans and a resolution of 8 cm^-1^. Background substruction and baseline correction were performed using Omnic Specta (Thermo Fischer), whereas plotting and spectrum analysis were performed using OriginPro 2020.

#### Scanning Electron Microscopy (SEM) Imaging

The mineral particles were mounted on an SEM stub using conductive carbon adhesive tape to determine their size and morphology. The particles were imaged using a field-emission SEM (Verois 460L, Thermo Fisher) at a voltage of 2-3 kV.

### 2.3 Cell culture and spheroid formation

MCF10DCIS.com immortalized human breast precancer epithelial cells were used (Barbara Ann Karmanos Cancer Institute). Cells were cultured in minimal DMEM/F12 media (1% Penicillin/Streptomycin, 5% horse serum) at 37 °C and 5% CO_2_ with refreshing media three times a week. The cells were routinely tested for mycoplasma contamination. For spheroid formation, we coated the bottom of each well in a 96-well plate with 70 μL of 1.5% low melting agarose in DMEM/F12 to prevent cell adhesion^35,38^. MCF10DCIS.com cells of 80% confluency were trypsinized and seeded at a density of 5000 cells/100 μL per well of minimal DMEM/F12 media and 30 μg/mL MC analogs (from one of the following minerals: COD, LCA, and HCA) into each well (Day 0). The minerals were ground with a pipette tip in 70% EtOH and sonicated for 15 min ahead to minimize their size and then sterilized with 70% EtOH. The minerals were dried in a biosafety cabinet under UV light for at least 20 min and until they were completely dry before adding them to the culture media. The 96-well plates were incubated at 37 °C and 5% CO_2_ on a shaker to encourage spheroid formation in suspension. On Day 1, 100 μL of minimal DMEM/F12 media were added. Every 48 h, 100 μL media were aspirated, and fresh 100 μL media were added to each well. The shaking and the low attachment coating ensured that only one spheroid formed in each well, resulting in a homogeneous spheroid population throughout the wells in the plate.

### 2.4 Spheroid imaging

#### 2.4.1 microCT imaging and analysis

Spheroids with and without LCA MC analogs were washed twice with phosphate-buffered saline (PBS) on Day 4 of the culture and fixed with 4% paraformaldehyde (PFA), n=5. The fixed spheroids were rewashed with PBS to eliminate any PFA in the sample. To increase the contrast of the organic material of the spheroids, they were stained with iodine by adding 0.5 mL deionized water, followed by 150 μL Lugol’s EMS solution (1% I_2_, 2% KI) for 30 min. The spheroids were washed with deionized water, transferred to 0.5 mL centrifuge tubes, and then mounted in a 2% low melting agarose solution at the bottom of the tube to enable scanning from all directions. The spheroids were scanned using a Zeiss Versa 520 microCT. Each scan was performed at 40kV and 5W settings, and the images were taken with a 10 s exposure time. The data were binned to a final resolution of 1.3–1.4 μm/pixel.

Image processing was performed using Avizo Fire software to 3D reconstruct the data and obtain 2D ortho slices and volume rendering. MC analogs and organic material, including cells and ECM, were outlined by thresholding, brush tools, and magic wand tools. The spheroids and MC analogs were segmented, and the spheroid diameters were measured using the Avizo Fire measurement tool.

#### 2.4.2 SEM and backscattered electron (BSE) spheroid analysis

Spheroids were collected on Day 7 of the culture into a 1.5 mL centrifuge tube, fixed with 4% PFA for 30 min, washed with PBS, and stained with Eosin Y solution 5% (G-Biosciences, St. Louis, USA) to enable localization during histological processing. Ten fixed spheroids were placed in a Tissue-Tek biopsy cryomold (Sakura) and embedded in an optimal cutting temperature compound (OCT, Scigen Scientific Gardena, CA, USA). Next, the molds were quickly put in liquid nitrogen, stored at −80 °C, and transferred to −20 °C 30 min before sectioning in a Leica CM3050S cryostat. Sectioning was performed at −22 °C, and 9-μm-thick sections were mounted on a glass slide covered with aluminum foil. The sections were kept on the aluminum foil glued to SEM stabs using PELCO Colloidal Graphite Isopropanol based (Ted Pella, Inc., USA) and then sputter-coated with carbon using a Quorum Q150T ES instrument. Imaging was carried out using a field-emission SEM (Verois 460L, Thermo Fisher) at a voltage of 3-10 kV, ETD (secondary electrons) and MD (backscattered electrons) detectors.

#### 2.4.3 Light microscopy imaging

The MC analogs were imaged immediately before they were added to the cells to evaluate their size while being dispersed in the culture media. After the MC analogs underwent the sterilization process, they were suspended in minimal DMEM/F12 media and were imaged using a Nikon ECLIPSE Ti2-U inverted microscope equipped with a DS-QI2 mono-cooled digital camera with a 40× objective.

Live unfixed spheroids were imaged using light microscopy to monitor their size and morphology. Spheroids were imaged on Days 1 and 7 of the culture using a Nikon ECLIPSE Ti2-U inverted microscope equipped with a DS-QI2 mono-cooled digital camera with a 10× objective. Adobe Photoshop was used for brightness and contrast adjustments. Image processing was performed consistently in all conditions. ImageJ software^39^ was used for spheroid diameter measurements. Spheroid diameters are presented as the mean ± standard deviation (n=9 for each condition).

### 2.5 Histopathology

Spheroid sections were prepared following the same procedure as the sections for SEM imaging; the sections were mounted on positively charged glass slides. The spheroid sections were stained with Hematoxylin stain, Gill no.2 (G-Biosciences, St. Louis, USA) for 5 seconds and Eosin Y (G-Biosciences, St. Louis, USA) for 1 second. Imaging was carried out using a Nikon Eclipse Ci-L Ergo R2S Fi3 microscope equipped with a DS-Fi3 camera with a 20× objective. Adobe Photoshop was used for brightness and contrast adjustments. Image processing was performed consistently in all conditions.

### 2.6 Cytotoxicity

The cytotoxicity of MC analogs and oxalate ions was studied in DCIS cell monolayers cultured in 96-well plates. For calibration, concentrations of 2500, 7500, and 10,000 cells were used. For cytotoxicity measurements, 5000 cells were seeded per well. They were cultured with 30 μg/mL of one of the following minerals: LCA, HCA, and COD or with 0.22M of sodium oxalate (to enrich the media with oxalate ions). This oxalate ion concentration was chosen because it is the maximum ion concentration that will be obtained if all the added COD crystals dissolve in the solution. The minerals were sterilized with 70% EtOH and UV light before being added to the culture media. The control cells were cultured without added minerals. After 24 h of incubation, the media was aspirated, and the cells were washed with PBS; then 50 μL of 0.1% crystal violet (CV) dissolved in deionized water were added to each well for 10 min. Next, the cells were washed with deionized water at least five times until the water became transparent. The plates were left to dry for at least 45 minutes and were imaged using a Nikon Eclipse Ci-L Ergo R2S Fi3 microscope equipped with a DS-Fi3 camera with a 4× objective. Next, 50 μL of acetic acid were added to each well to dissolve the CV, and the absorbance at 570 nm was measured with an Infinite M200 plate reader (TECAN).

The experiment was repeated with two independent biological replicates using six technical replicates each. The absorbance of the calibration samples was used to create the linear calibration curve, from which the calibration equation that correlates between absorbance and the cell number was extracted using OriginPro 2020. This calibration equation was used to determine the cell number in each sample.

### 2.7 Western blotting

The expression of Her2 was studied in 3D multicellular spheroids. Thirty spheroids from each condition were collected into a 1.5 mL centrifuge tube, washed with PBS, and centrifuged for 5 min at 5 rpm and at 4°C. The spheroids were lysed by adding 100 μL of ice-cold RIPA buffer (50 mM Tris (Sigma Aldrich), 150 mM NaCl (Sigma Aldrich), 0.5% Triton (Sigma Aldrich), 0.1% SDS (Sigma Aldrich) containing 10% protease inhibitor cocktail (Sigma Aldrich) and incubated on ice for 40 min. The cell debris was pelleted by centrifugation for 10 min at 15,000 rpm and at 4°C, and the supernatant was collected. The protein concentration of the lysates was determined using the BCA protein assay (Pierce). The samples for Western blotting were prepared by mixing 15 μg protein from each spheroid lysate with Laemmli Sample Buffer, boiling it for 5 min, and then chilling on ice. The proteins were separated by electrophoresis on 10% agarose gel and transferred to a nitrocellulose membrane. The membrane was incubated for 1 h in a blocking buffer of PBST (PBS (Sigma Aldrich) supplemented with 0.1 v/v % Tween 20 (Sigma Aldrich) and 5% (v/w) dry skim milk (Sigma-Aldrich). The membrane was washed three times with PBST for 5 min and then incubated with primary antibody diluted in PBST supplemented with 2% skim milk. The anti-Her2 antibody (sc-33684, Santa Cruz Biotechnology, Dallas, TX, USA) was diluted 1:50 (v/v) in PBST supplemented with 2% skim milk prior to use, and incubated with the membrane for 3h. The anti β-Actin antibody (sc-47778) was diluted 1:400 (v/v) and incubated with the membrane for 1 h. Next, the membrane was washed three times with PBST and incubated for 1h at 25°C with a secondary, anti-mouse HRP-conjugated antibody (115-035-003, Jackson ImmunoResearch Laboratories, West Grove, PA, USA) diluted 1:10,000 (v/v) in PBST supplemented with 0.5% (v/w) skim milk. To finalize the procedure, the membrane was washed three times with PBST and incubated with an ECL Western blotting reagent (1,705,060, Bio-Rad, Hercules, CA, USA) for 5 min in the dark. Images were collected by chemiluminescence, using the Fusion FX imaging system (Vilber Lourmat, Collégien, France).

### 2.8 Statistical analysis

All data were expressed as the mean +/- standard deviation (SD). One-way ANOVA analysis was used to compare differences between the diameter and the cytotoxicity experiments. P-values less than 0.05 were considered significant. *P < 0.05. Statistical analysis was performed using OriginPro 2020. All experiments were performed with at least two independent biological replicates, each with at least three technical replicates.

## 3. Results

### 3.1 Fabrication of spheroids containing synthetic MC analogs

We developed an engineered 3D breast precancer tumor model in which synthetic mineral particles with the desired crystal phase and properties serve as MC analogs.

The synthetic MC analogs’ low and highly carbonated apatite and COD were ground and sonicated in 70% ethanol and underwent additional sterilization with UV light. MCF10DCIS.com precancer cells were mixed with the MC analogs in a 96-well plate under low attachment conditions, leading to the formation of one multicellular spheroid in each well (Fig. 1). To determine how varying the composition and crystal properties of MC analogs affects the precancer cell behavior, only one type of MC analog was introduced to each multicellular spheroid. The effect of the mineral type on cell behavior was studied after seven days of culture because spheroids cultured for extended time periods spontaneously deposit hydroxyapatite mineral particles^35^.

**Figure 1.**
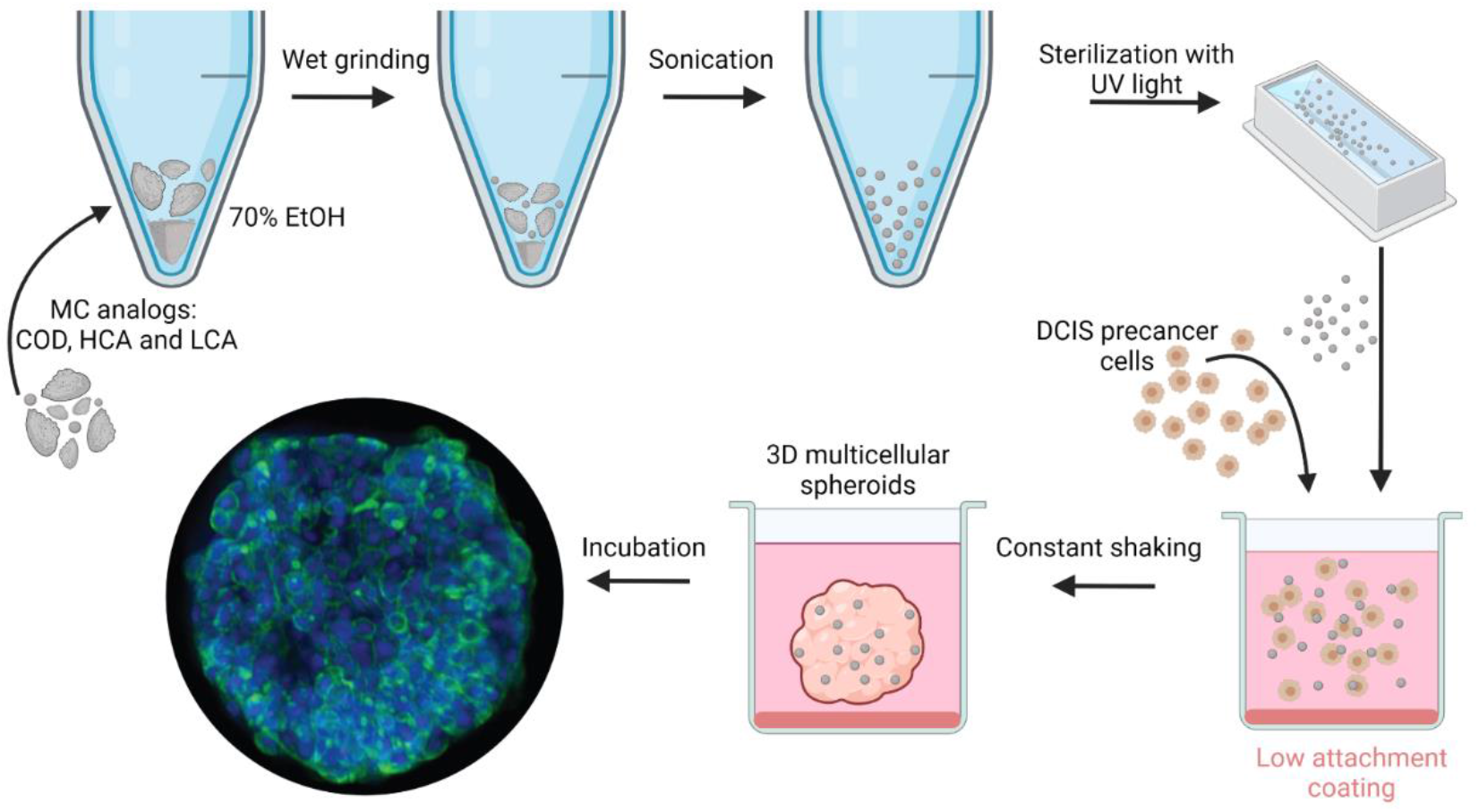
The experimental design for creating precancer multicellular spheroids containing MC analogs. The MC analogs were ground in 70% EtOH and then sonicated to minimize their size. Then, they were sterilized with ethanol and UV light. The sterile and dry MC analogs were added into 96-well plates with a cell dispersion of 50,000 cells/mL, and the plates were constantly shaken. After incubation, 3D spheroids form in suspension, and the effect of MC analogs on precancer cells can be measured. The MC analogs used are COD, HCA, and LCA. Spheroids without added minerals served as the control.

We used commercial LCA and in-house synthesized HCA and COD crystals for MC analogs. The LCA, HCA, and COD crystals differ in their composition, crystal phase, morphology, particle size, and surface texture. We determined the crystal phase of LCA, HCA, and COD by Fourier Transform InfraRed spectroscopy (FTIR) (Fig. 2 A). The v_1_ and v_3_ phosphate peaks were observed in LCA and HCA, and for LCA, the v_1_ OH peak was observed as well, indicating that it resembles stoichiometric hydroxyapatite. The v_2_ carbonate peaks were observed for LCA and HCA, and the intensity of the v_3_ carbonate peak normalized to the v_3_ phosphate peak was lower in LCA, compared with HCA. The carbonate/phosphate peak area ratio was 0.009 for HCA and 0.008 for LCA, and the LCA has a higher crystallinity than does HCA, as indicated by its phosphate peak full width at half of its maximum height (70 cm^-1^ for LCA and 107 cm^-1^ for HCA). Carbonyl peaks (C=O and C-O stretch) were observed in the COD spectra.

**Figure 2.**
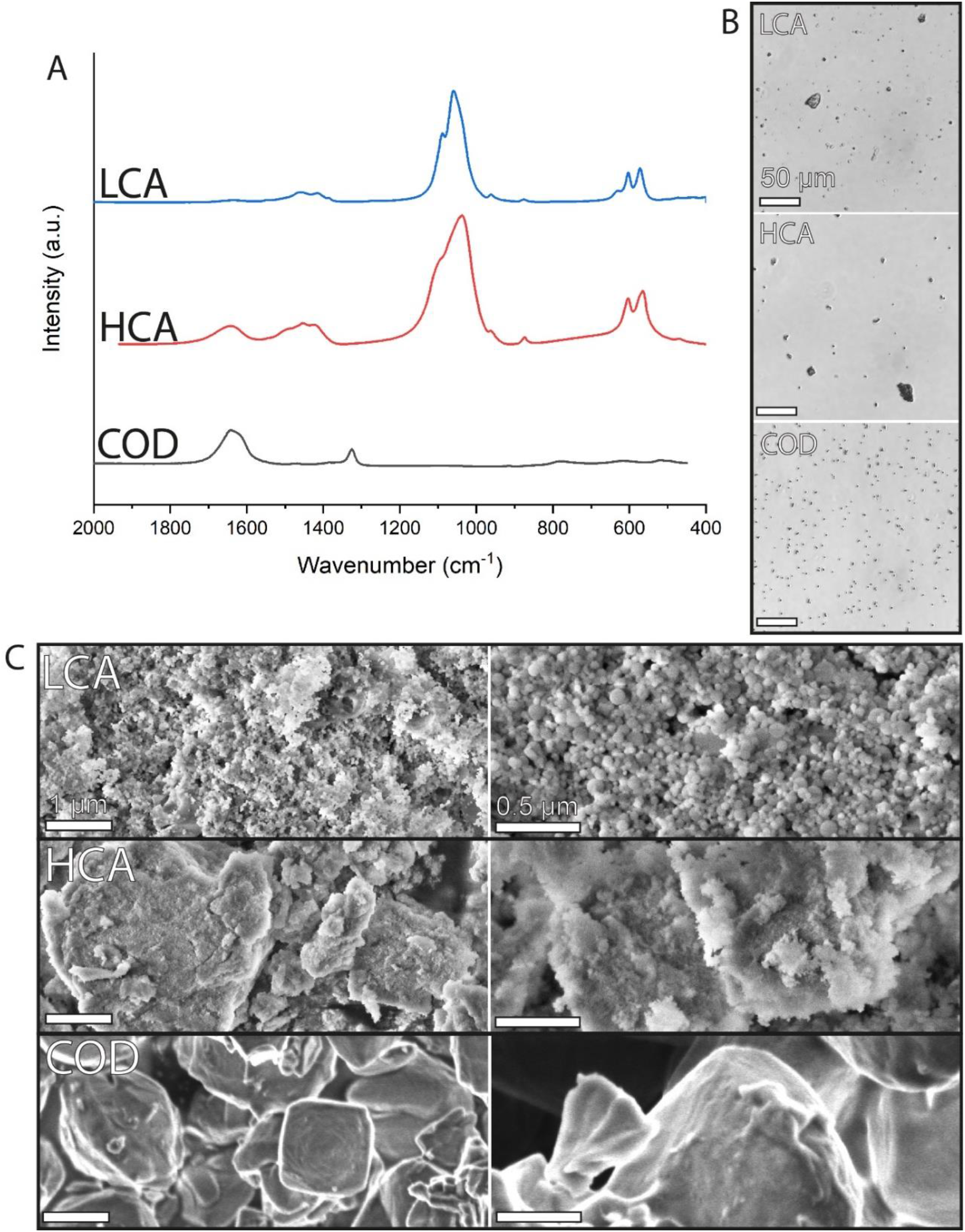
Characterization of the MC analogs LCA, HCA, and COD before they are added to the cells to form multicellular spheroids. **A.** FTIR spectra of the crystals. **B.** Light microscope images of the crystals suspended in culture media. Sacle bars are 50 μm. **C.** SEM micrographs of the crystals showing the crystal morphology, particle sizes, and surface texture. Scale bars – left: 1 μm, right: 0.5 μm.

Each MC analog disperses differently when added to the culture media (Fig. 2 B). HCA and LCA particles aggregate to form particles in the size range of 1.5-30 μm and 1.1-20 μm, respectively. COD particles, however, do not aggregate, are homogeneously dispersed in the culture media, and are in the size range of 1.5-3 μm. These changes in the particle’s tendency to aggregate most likely result from differences in the crystal surface properties.

LCA, HCA, and COD crystals also differ in their morphology, surface texture, and particle size, as observed by SEM imaging (Fig. 2 C). The LCA crystals are sub-micrometer spherical particles loosely aggregated together, whereas the HCA crystals are denser and larger. The COD crystals are smaller than 3 μm, with a smooth surface, some showing typical bipyramidal morphology and others with less clear facets. These properties most likely play a role in crystal-crystal, crystal-protein, and crystal-cell interactions.

We used suspensions of varying mineral concentrations in culture media to develop the protocol for embedding the synthetic MC analogs in precancer spheroids. We monitored the spheroid culture using light microscopy to optimize the conditions in which the mineral is incorporated into the spheroid. After choosing an optimal mineral concentration of 30 μg/mL, we further verified that the MC analogs are indeed embedded within the spheroids and that their incorporation did not disrupt the spheroid formation. Because spheroids consist of dense organic material and have diameters of hundreds of micrometers, light microscopy imaging was insufficient to characterize their interior. We compared spheroids with and without the MC analog LCA using microCT 3D X-Ray imaging (Fig. 3 A-D), which allows for obtaining 3D images of the interior of spheroids. The cellular and extracellular organic material, the inorganic MC analogs, and lumens were all observed in the spheroids. They were distinguished based on their density, which produces distinct contrast in the microCT images. MC analogs, which are calcium-containing crystals, are observed in the microCT images as bright particles due to their high atomic number compared with their surroundings. MC analogs were not observed in control spheroids (Fig. 3 A, C). In contrast, MCs were observed in spheroids to which synthetic MC analogs were added (Fig. 3 B, D), validating that only the precancer spheroids contain mineral particles. The 2D ortho slices of the spheroids show that the observed particles are distributed within the spheroid volume and are not only concentrated in specific areas (Fig. 3 B), most likely leading to many cell-mineral interactions. The sizes of the observed MC analogs reach up to 75 μm, which is much larger than their sizes in the suspension. The embedded mineral particles can be characterized in situ and in high resolution using SEM imaging of the spheroid sections (Fig. 3 F, Fig. S1). Owing to their high atomic number and elemental composition, compared with their organic surroundings, the mineral particles can be identified as brighter features in the sections via backscattered electron detection (Fig. S1). In agreement with the microCT results, the mineral particles were only observed by SEM imaging in spheroids with added MC analogs. Energy-dispersive X-Ray spectroscopy (EDS) verified that the elemental composition of the particles is as expected (Fig. S1). Spheroids with LCA MC analogs are larger than those without the MC analogs. Lumens that spontaneously form in the spheroid structure are observed in the spheroid microCT and SEM images.

**Figure 3.**
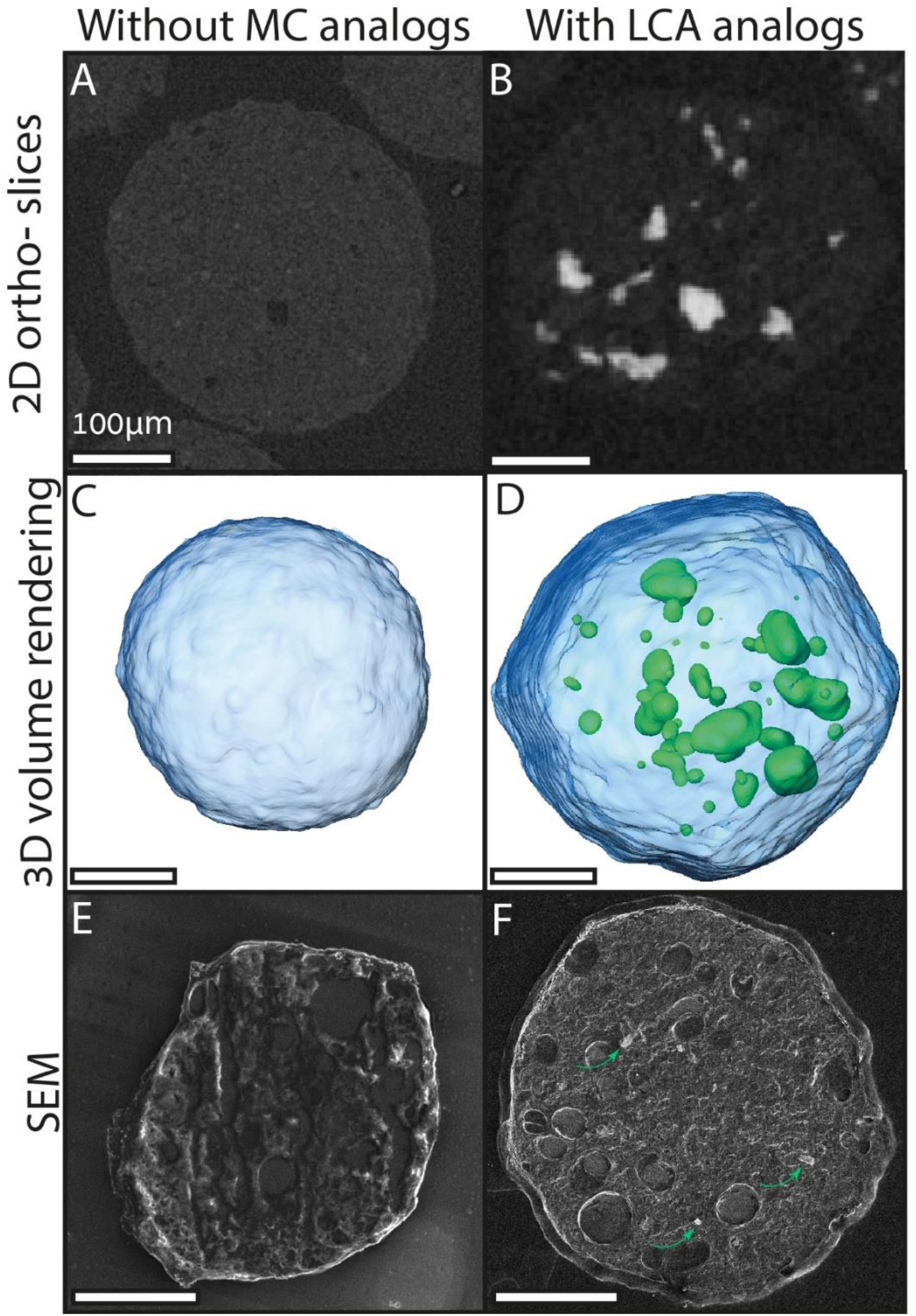
Characterization of multicellular spheroids with and without added LCA MC analogs. **A-D**. microCT 3D X-ray imaging of spheroids on Day 4 of the culture. **A, B**. 2D ortho slices from 3D reconstructed spheroids showing the spheroid center, where mineral particles of high atomic number, compared with the organic surrounding, are observed as white features. **C, D**. Volume rendering of the spheroids showing the mineral particles embedded within the spheroids, where the MC analogs are denoted in green and the organic material is in blue. **E, F**. SEM images of sectioned spheroids on Day 7 of the culture. **F.** Small particles that correspond to a backscattered electron (BSE) signal are observed in the spheroid section (green arrows). All sacle bars are 100 μm.

### 3.2 Spheroid diameter and morphology according to the MC analog type

To study the effect of the mineral type on precancer spheroid morphology and diameter, spheroids with added MC analogs and control spheroids without added minerals were imaged on Day 1 and Day 7 of the culture (Fig. 4). To characterize the spheroid interior further, histopathological sections were taken from each spheroid, stained with Hematoxylin and Eosin (H&E), and imaged under cross-polarized light to identify the birefringent particles (Fig. 4).

**Figure 4.**
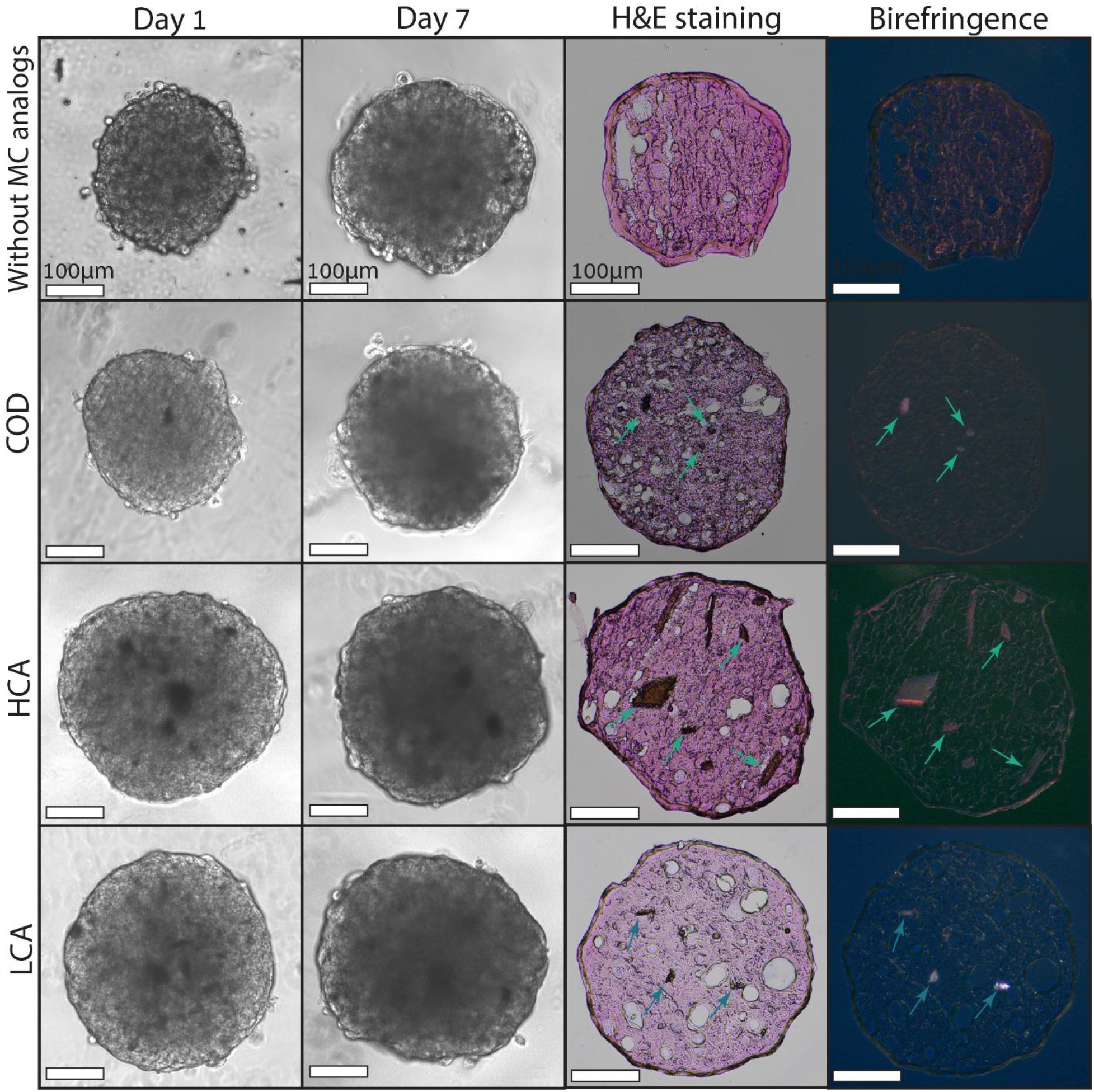
Light microscopy images of precancer spheroids with and without embedded MC analogs. **Left:** Live spheroids on Days 1 and 7 of the culture. **Right:** Histopathology of spheroid sections after seven days of culture, taken with brightfield and cross-polarized light. MC analogs embedded within the tissue sections can be observed (green arrows). All sacle bars are 100 μm.

On Day 1 of the culture, a difference in diameter was already observed between spheroids according to the MC analog mineral type. Spheroids with calcium phosphate particles were larger than the control spheroids and spheroids with COD. The spheroids became less transparent between Day 1 and Day 7 of the culture for all MC analog types. This decrease in spheroid transparency suggests that the spheroids become denser with time, most likely due to cell proliferation and extracellular matrix secretion. The dark features dispersed in the spheroids are most likely materials that differ from the cells, possibly the MC analogs or lipid droplets. On Day 7, spheroids without added MC analogs and those containing COD had the smallest diameter, followed by spheroids with HCA. Spheroids with LCA had the largest diameter (Table 1).

**Table 1.** Average spheroid diameter according to the culture time and MC analog type

|  | Day 1 | P-value | Day 7 | P-value |
| --- | --- | --- | --- | --- |
| **Without MC analogs** | $320 \pm 40 \mu\text{m}$ | | $340 \pm 10 \mu\text{m}$ | |
| **COD MC analogs** | $340 \pm 30 \mu\text{m}$ | 0.415 | $350 \pm 20 \mu\text{m}$ | 0.218 |
| **HCA MC analogs** | $380 \pm 20 \mu\text{m}$ | <0.05 | $400 \pm 20 \mu\text{m}$ | <0.05 |
| **LCA MC analogs** | $380 \pm 10 \mu\text{m}$ | <0.05 | $400 \pm 20 \mu\text{m}$ | <0.05 |

### 3.3 Her2 expression in precancer spheroids according to the MC analog type

We assessed the expression of Her2, associated with cancer cell growth, in response to 7 days of culture with varying MC analog types, using Western blot (Fig. 5 A, Fig. S2). Precancer cells cultured in spheroids with the calcium phosphate MC analogs LCA and HCA show an increased expression of Her2 compared with spheroids with no MC analogs. However, cancer cells cultured with COD show decreased Her2 expression compared with calcium phosphate-containing spheroids and spheroids with no added mineral particles.

**Figure 5.**
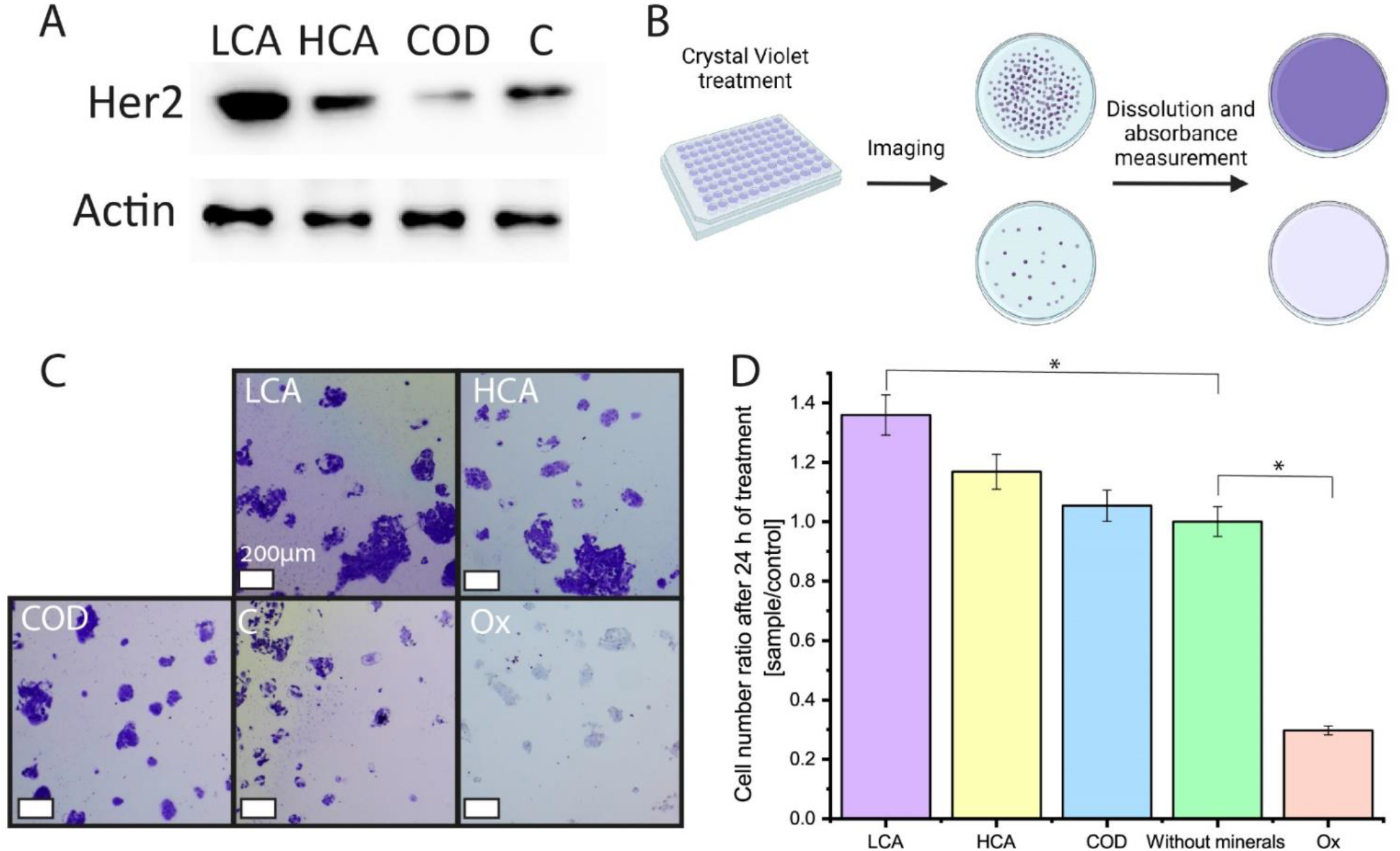
The effect of MC analogs on DCIS cell behavior. **A.** WB analysis for Her2 expression in spheroids on Day 7 of the culture according to the MC analog type. **B.** Cytotoxicity assessment protocol. **C.** DCIS cell monolayers stained with Crystal Violet (CV) after treatment with different MC analogs for 24h. **D.** The effect of MC analogs and dissolved oxalate ions on cell viability was measured by a CV assay after 24 h of treatment. Cell numbers were normalized to the control. Errors represent the standard deviation. C = control, Ox = oxalate ions dissolved in the culture media.

### 3.4 Cytotoxicity of MC analogs

The decrease in Her2 expression in COD-containing spheroids, compared with control spheroids, suggests that COD has a reducing effect on the progression of DCIS cells. To determine whether the COD mineral, rather than its *in-situ* dissolution into oxalate ions, induces changes in DCIS cell behavior, we compared the cell proliferation in response to treatment with MC analogs, including COD minerals and oxalate ions dissolved in the culture media. The concentration of oxalate ions was calculated as if the embedded mineral particles dissolved entirely. After LCA and HCA treatment, more cells are visible on the plates compared with cells treated with COD or control cells (Fig. 5 C). After treatment with culture media enriched with oxalate ions, very few live cells were detected on the plates. Quantitively, although the cell number after COD treatment was similar to the control, there was an increase in the cell number as a response to treatment with calcium phosphate mineral particles, where a larger cell number was obtained after treatment with LCA. The cell number dropped after oxalate ion treatment, indicating that exposure to dissolved oxalate ions harms the cells (Fig. 5 D). This drop suggests that it is unlikely that the precancer cell response to COD is due to its dissolution to ions, since the effect on the cells of COD minerals and oxalate ions is very different. Since COD may induce tumor suppression and potentially be used to treat precancer tumors, this proliferation test also assesses the treatment cytotoxicity, validating that COD is not cytotoxic.

## 4. Discussion

MCs are a component of the breast tumor microenvironment, and their chemical and physical properties correlate with lesion malignancy. However, it is not clear whether the MC properties affect disease progression, whether cancer-induced changes in the tissue lead to these properties, or whether there is some combination of both. *In-vitro* and *in-vivo* studies indicate that apatite crystals can trigger cancer behavior and that the cellular response to the apatite depends on its properties. The effect of hydroxyapatite crystals on normal or precancer breast cells was examined in cell monolayers or cells cultured in mineral-containing scaffolds but never in a 3D tumor model such as multicellular spheroids. These *in-vitro* studies showed increased mitogenesis, proliferation, cell adhesion, matrix metalloproteinase upregulation, and secretion of pro-osteoclastic interleukin-8 (IL-8) as a response to hydroxyapatite crystals^10,22–25^. Additionally, *in-vitro* studies examining the effect of minerals on cancer progression focused on calcium phosphate minerals such as hydroxyapatite, assuming that the calcium oxalate mineral has no effect on cells and serves as a bystander in the tissue microenvironment because it is commonly associated with benign lesions.

We developed a DCIS 3D tumor model with embedded synthetic MC analogs that are physiologically relevant and whose composition and crystal properties can be controlled. We showed that precancer cells cultured as 2D monolayers and as 3D multicellular spheroids mimicking DCIS tumors respond differently to varying MC analog types. When DCIS cells encounter LCA crystals in their environment, and to a lesser extent also HCA crystals, their spheroid diameter and Her2 expression increase. When DCIS cells encounter COD crystals, their spheroid diameters resemble those of control cells, and are smaller than those of LCA and HCA, whereas Her2 expression is less than in control cells. These malignancy-associated changes in cell behavior indicate that MC properties can affect disease progression, since apatite crystals affect Her2 expression according to the carbonate fraction, whereas COD crystals suppress Her2 expression. Her2 is a transmembrane receptor that is part of the epidermal growth factor family, which regulates cell growth and proliferation^40^. Overexpression of Her2 occurs in 15-30% of invasive breast cancers^40,41^ and correlates with more aggressive tumor behavior, a poorer prognosis, and increased mortality^42,43^. Since Her2 overexpression is a marker of poor prognosis, the changes in Her2 expression reported here suggest that COD has a suppressive effect on the precancer cells. In contrast, HCA and LCA trigger cancerous behavior.

In our model, the DCIS cells are cultured in multicellular spheroids that contain MC analogs, and not all cells are in direct contact with the MC analogs, much like in actual tumors. Since the culture media is supersaturated with respect to apatite and COD crystals, which have low Ksp values, we assumed that the effect of these crystals on the cells is due to physical contact and not via ion dissolution. To verify this, we added oxalate ions to the culture media and showed that COD crystals affect the cells differently than do soluble oxalate ions, which harmed the cells. A possible reason for this harmful effect is that oxalate ions can activate signaling pathways, DNA synthesis, cell growth, IL-7 production, and are known to have a toxic effect that can lead to apoptosis in renal epithelial cells^44,45^. However, when breast cancer cells were exposed to oxalate ions *in vitro*, their proliferation and the expression of a pro-tumorigenic gene increased^46^. The difference between our observation of decreased cell proliferation, when exposed to oxalate ions, and the abovementioned observation of increased proliferation, may result from the higher oxalate concentrations that we used (0.22 mM vs. 20-50 μM) or the difference in cell type.

Spheroids with apatite are larger than those with COD, even on Day 1 of the culture. Some possible explanations are that synthetic apatite crystals induce deposition of apatite crystals by the cells or via heterogeneous nucleation and seeded growth, increasing the total crystal volume and consequently the spheroid size. This is in agreement with the observation that LCA particles within the spheroids were larger than the ones dispersed in the culture media before being added to the cells. Another possibility is that apatite crystals trigger precancer proliferation already in the first 24 h of culture, and the higher cell number is the reason for the larger spheroids. Additionally, the cells in spheroids with COD may be more tightly/densely packed together than the cells in spheroids with apatite because they remain more epithelial than the potentially more mesenchymal motile cells in contact with apatite.

The MC analogs used in this study were chosen based on their crystal phase and composition because in clinical samples of breast biopsies malignancy is correlated with the MC crystal phase and the composition. MCs and MC analogs, however, differ not only in their chemistry; they also differ in their particle size, crystal morphology, surface area, and roughness^26^. Here, the LCA crystals consisted of smooth sub-micron particles, whereas the HCA crystals had larger particle sizes and rougher surfaces. The COD crystals used were smooth, with sizes up to 3 μm. Whereas the LCA and HCA crystals aggregated in the culture media, the COD crystals exhibited a low tendency to aggregate. Most likely, the smaller particle size and the high surface topography of LCA and HCA induce protein adsorption that increases aggregation, unlike the smoother COD surface. This aggregation can be mediated by proteins, since protein adsorption to mineral surfaces depends on the protein conformation, the crystal particle sizes, the surface topography, roughness, porosity, pore size, charge, and the functional groups^47^.

Most likely, these MC analog characteristics play a role in the mineral-cell interactions inside the spheroid. Differences in the morphology, surface roughness, and area possibly translate to changes in the mineral’s tendency to adhere to cells and aggregate^48^. Such differences in mineral-cell interactions can either promote or inhibit the aggregation of cells and minerals and induce signaling pathways leading to more malignant cell behavior. For example, COD and calcium oxalate monohydrate (COM) are both calcium oxalate crystals found in kidney stones. In the context of the renal system, the structure and composition of these minerals vary and result in differences in the ability of epithelial cells to attach to these crystals, favoring COM attachment. This was suggested to be why COM has a much more harmful effect on the renal system than COD does, because it triggers crystal aggregation *in vitro* and may lead to changes in the epithelial characteristics of cells and consequently transform them into more mesenchymal cells^49,50^. Furthermore, calcium oxalate crystal shape and aggregation affected the renal epithelial cells: the cells adhered more to crystals with large calcium ion-rich faces, which exhibited the largest cytotoxicity^51^. Renal cells exposed to bipyramidal COD crystals similar in shape and size to the crystals in our study exhibited a high viability^52^. Future studies may also consider MC-cell interactions in DCIS, including mixing MC analogs, the co-culture of cells, monitoring crystal growth and aggregation in the spheroid, as well as elucidating the underlying biomolecular mechanisms related to cell-mineral attachment. Taken together, COD, which is often found in benign lesions, is possibly not just a bystander but instead, it locally suppresses DCIS cells in its proximity; hence, it is rarely found in invasive tissues.

## 5. Conclusions

We developed a model of 3D precancer spheroids consisting of synthetic MC analogs with mineral properties that can be rationally designed and controlled. As reflected by their spheroid diameter and Her2 expression, the malignancy potential of breast precancer cells changes with the mineral properties of the embedded MC analogs in our *in-vitro* model. LCA promoted precancer malignancy compared with other minerals or conditions in which no mineral was added. COD suppresses precancer malignancy compared with apatite and decreases Her2 expression compared with apatite and the control. Unlike dissolved oxalate ions, COD is not cytotoxic. Potentially, insights from this study may provide new directions to inhibit the progression of DCIS precancerous lesions to invasive breast cancer using tumor-suppressing mineral particles and to evaluate DCIS prognosis based on COD as a biomarker.

## Author contributions

AC designed and performed the study, analyzed and visualized the results, and wrote the initial draft of the manuscript. LG and DA contributed to the methodology and analysis and reviewed the final draft of the manuscript. NV conceptualized, designed, and supervised the study, as well as reviewed and edited the manuscript.

## Conflicts of interest

The authors declare no conflicts of interest.

## Acknowledgments

This work was supported by the Israeli Science Foundation [grant number 565/21]. NV is the incumbent of the Joseph and May Winston Career Development Chair in Chemical Engineering. The authors thank Vlad Brumfeld and Sergey Kaphishnikov for their assistance with microCT measurements and Roxana Golan and Sahar Koresh for their help with the SEM imaging. The illustrations were created with BioRender.com.

## Notes

### Competing Interest Statement

The authors have declared no competing interest.

## References

(1) Nelson, A. C.; Machado, H. L.; Schwertfeger, K. L. Breaking through to the Other Side: Microenvironment Contributions to DCIS Initiation and Progression. J. Mammary Gland Biol. Neoplasia 2018, 23 (4), 207–221. https://doi.org/10.1007/s10911-018-9409-z.

(2) Hofvind, S.; Iversen, B. F.; Eriksen, L.; Styr, B. M.; Kjellevold, K.; Kurz, K. D. Mammographic Morphology and Distribution of Calcifications in Ductal Carcinoma in Situ Diagnosed in Organized Screening. Acta radiol. 2011, 52 (5), 481–487. https://doi.org/10.1258/ar.2011.100357.

(3) Holland, R.; Hendriks, J. H. C. L. Microcalcifications Associated with Ductal Carcinoma in Situ: Mammographic-Pathologic Correlation. Semin. Diagn. Pathol. 1994, 11 (3), 181–192.

(4) Tabar, L.; Chen, H. H. T.; Yen, M. F. A.; Tot, T.; Tung, T. H.; Chen, L. S.; Chiu, Y. H.; Duffy, S. W.; Smith, R. A. Mammographic Tumor Features Can Predict Long-Term Outcomes Reliably in Women with 1-14-Mm Invasive Breast Carcinoma: Suggestions for the Reconsideration of Current Therapeutic Practice and the TNM Classification System. Cancer 2004, 101 (8), 1745–1759. https://doi.org/10.1002/cncr.20582.

(5) Thurfjell, E.; Thurfjell, M. G.; Lindgren, A. Mammographic Finding as Predictor of Survival in 1-9 Mm Invasive Breast Cancers. Worse Prognosis for Cases Presenting as Calcifications Alone. Breast Cancer Res. Treat. 2001, 67 (2), 177–180. https://doi.org/10.1023/A:1010648919150.

(6) Rauch, G. M.; Hobbs, B. P.; Kuerer, H. M.; Scoggins, M. E.; Benveniste, A. P.; Park, Y. M.; Caudle, A. S.; Fox, P. S.; Smith, B. D.; Adrada, B. E.; Krishnamurthy, S.; Yang, W. T. Microcalcifications in 1657 Patients with Pure Ductal Carcinoma in Situ of the Breast: Correlation with Clinical, Histopathologic, Biologic Features, and Local Recurrence. Ann. Surg. Oncol. 2016, 23 (2), 482–489. https://doi.org/10.1245/s10434-015-4876-6.

(7) Elias, S. G.; Adams, A.; Wisner, D. J.; Esserman, L. J.; Van’t Veer, L. J.; Mali, W. P. T. M.; Gilhuijs, K. G. A.; Hylton, N. M. Imaging Features of HER2 Overexpression in Breast Cancer: A Systematic Review and Meta-Analysis. Cancer Epidemiol. Biomarkers Prev. 2014, 23 (8), 1464–1483. https://doi.org/10.1158/1055-9965.EPI-13-1170.

(8) Kunitake, J. A. M. R.; Choi, S.; Nguyen, K. X.; Lee, M. M.; He, F.; Sudilovsky, D.; Morris, P. G.; Jochelson, M. S.; Hudis, C. A.; Muller, D. A.; Fratzl, P.; Fischbach, C.; Masic, A.; Estroff, L. A. Correlative Imaging Reveals Physiochemical Heterogeneity of Microcalcifications in Human Breast Carcinomas. J. Struct. Biol. 2018, 202 (1), 25–34. https://doi.org/10.1016/j.jsb.2017.12.002.

(9) Vidavsky, N.; Kunitake, J. A. M. R.; Estroff, L. A. Multiple Pathways for Pathological Calcification in the Human Body. Adv. Healthc. Mater. 2020, 2001271 (4), 1–23. https://doi.org/10.1002/adhm.202001271.

(10) Cox, R. F.; Hernandez-Santana, A.; Ramdass, S.; McMahon, G.; Harmey, J. H.; Morgan, M. P. Microcalcifications in Breast Cancer: Novel Insights into the Molecular Mechanism and Functional Consequence of Mammary Mineralisation. Br. J. Cancer 2012, 106 (3), 525–537. https://doi.org/10.1038/bjc.2011.583.

(11) Baker, R.; Rogers, K. D.; Shepherd, N.; Stone, N. New Relationships between Breast Microcalcifications and Cancer. Br. J. Cancer 2010, 103 (7), 1034–1039. https://doi.org/10.1038/sj.bjc.6605873.

(12) Frappart, L.; Boudeulle, M.; Boumendil, J.; Lin, H. C.; Martinon, I.; Palayer, C.; Mallet-Guy, Y.; Raudrant, D.; Bremond, A.; Rochet, Y.; Feroldi, J. Structure and Composition of Microcalcifications in Benign and Malignant Lesions of the Breast: Study by Light Microscopy, Transmission and Scanning Electron Microscopy, Microprobe Analysis, and X-Ray Diffraction. Hum. Pathol. 1984, 15 (9), 880–889. https://doi.org/10.1016/S0046-8177(84)80150-1.

(13) Scott, R.; Stone, N.; Kendall, C.; Geraki, K.; Rogers, K. Relationships between Pathology and Crystal Structure in Breast Calcifications: An in Situ X-Ray Diffraction Study in Histological Sections. npj Breast Cancer 2016, 2 (1), 16029. https://doi.org/10.1038/npjbcancer.2016.29.

(14) Haka, A. S.; Shafer-Peltier, K. E.; Fitzmaurice, M.; Crowe, J.; Dasari, R. R.; Feld, M. S. Identifying Microcalcifications in Benign and Malignant Breast Lesions by Probing Differences in Their Chemical Composition Using Raman Spectroscopy. Cancer Res. 2002, 62 (18), 5375–5380.

(15) C J D’Orsi, F R Reale, M A Davis, V. J. B. Is Calcium Oxalate an Adequate Explanation for Nonvisualization of Breast Specimen Calcifications? RSNA Radiol. 1992, 182.

(16) Scimeca, M.; Giannini, E.; Antonacci, C.; Pistolese, C. A.; Spagnoli, L. G.; Bonanno, E. Microcalcifications in Breast Cancer: An Active Phenomenon Mediated by Epithelial Cells with Mesenchymal Characteristics. BMC Cancer 2014, 14 (1). https://doi.org/10.1186/1471-2407-14-286.

(17) Winston, J. S.; Yeh, I. T.; Evers, K.; Friedman, A. K. Calcium Oxalate Is Associated with Benign Breast Tissue: Can We Avoid Biopsy? Am. J. Clin. Pathol. 1993, 100 (5), 488–492. https://doi.org/10.1093/ajcp/100.5.488.

(18) Singh, N.; Theaker, J. M. Calcium Oxalate Crystals (Weddellite) within the Secretions of Ductal Carcinoma in Situ - A Rare Phenomenon. J. Clin. Pathol. 1999, 52 (2), 145–146. https://doi.org/10.1136/jcp.52.2.145.

(19) Martin, H. M.; Bateman, A. C.; Theaker, J. M. Calcium Oxalate (Weddellite) Crystals within Ductal Carcinoma in Situ [2]. J. Clin. Pathol. 1999, 52 (12), 932. https://doi.org/10.1136/jcp.52.12.932.

(20) Tornos, C.; Silva, E.; El-Naggar, A.; Pritzker, K. P. H. Calcium Oxalate Crystals in Breast Biopsies: The Missing Microcalcifications. Am. J. Surg. Pathol. 1990, 14 (10), 961–968. https://doi.org/10.1097/00000478-199010000-00010.

(21) O’Grady, S.; Morgan, M. P. Microcalcifications in Breast Cancer: From Pathophysiology to Diagnosis and Prognosis. Biochim. Biophys. Acta - Rev. Cancer 2018, 1869 (2), 310–320. https://doi.org/10.1016/j.bbcan.2018.04.006.

(22) Morgan, M. P.; Cooke, M. M.; Christopherson, P. A.; Westfall, P. R.; McCarthy, G. M. Calcium Hydroxyapatite Promotes Mitogenesis and Matrix Metalloproteinase Expression in Human Breast Cancer Cell Lines. Mol. Carcinog. 2001, 32 (3), 111–117. https://doi.org/10.1002/mc.1070.

(23) Pathi, S. P.; Kowalczewski, C.; Tadipatri, R.; Fischbach, C. A Novel 3-D Mineralized Tumor Model to Study Breast Cancer Bone Metastasis. PLoS One 2010, 5 (1), 1–10. https://doi.org/10.1371/journal.pone.0008849.

(24) Pathi, S. P.; Lin, D. D. W.; Dorvee, J. R.; Estroff, L. A.; Fischbach, C. Hydroxyapatite Nanoparticle-Containing Scaffolds for the Study of Breast Cancer Bone Metastasis. Biomaterials 2011, 32 (22), 5112–5122. https://doi.org/10.1016/j.biomaterials.2011.03.055.

(25) He, F.; Springer, N. L.; Whitman, M. A.; Pathi, S. P.; Lee, Y.; Mohanan, S.; Marcott, S.; Chiou, A. E.; Blank, B. S.; Iyengar, N.; Morris, P. G.; Jochelson, M.; Hudis, C. A.; Shah, P.; Kunitake, J. A. M. R.; Estroff, L. A.; Lammerding, J.; Fischbach, C. Hydroxyapatite Mineral Enhances Malignant Potential in a Tissue-Engineered Model of Ductal Carcinoma in Situ (DCIS). Biomaterials 2019, 224, 119489. https://doi.org/10.1016/j.biomaterials.2019.119489.

(26) Choi, S.; Coonrod, S.; Estroff, L.; Fischbach, C. Chemical and Physical Properties of Carbonated Hydroxyapatite Affect Breast Cancer Cell Behavior. Acta Biomater. 2015, 24, 333–342. https://doi.org/10.1016/j.actbio.2015.06.001.

(27) Jong, B. K.; Stein, R.; O’Hare, M. J. Three-Dimensional in Vitro Tissue Culture Models of Breast Cancer - A Review. Breast Cancer Res. Treat. 2004, 85 (3), 281–291. https://doi.org/10.1023/B:BREA.0000025418.88785.2b.

(28) Infanger, D. W.; Lynch, M. E.; Fischbach, C. Engineered Culture Models for Studies of Tumor-Microenvironment Interactions. Annu. Rev. Biomed. Eng. 2013, 15 (1), 29–53. https://doi.org/10.1146/annurev-bioeng-071811-150028.

(29) Edmondson, R.; Broglie, J. J.; Adcock, A. F.; Yang, L. Three-Dimensional Cell Culture Systems and Their Applications in Drug Discovery and Cell-Based Biosensors. Assay Drug Dev. Technol. 2014, 12 (4), 207–218. https://doi.org/10.1089/adt.2014.573.

(30) Breslin, S.; O’Driscoll, L. The Relevance of Using 3D Cell Cultures, in Addition to 2D Monolayer Cultures, When Evaluating Breast Cancer Drug Sensitivity and Resistance. Oncotarget 2016, 7 (29), 45745–45756. https://doi.org/10.18632/oncotarget.9935.

(31) Vidavsky, N.; Kunitake, J. A. M. R.; Diaz-Rubio, M. E.; Chiou, A. E.; Loh, H. C.; Zhang, S.; Masic, A.; Fischbach, C.; Estroff, L. A. Mapping and Profiling Lipid Distribution in a 3D Model of Breast Cancer Progression. ACS Cent. Sci. 2019, 5 (5), 768–780. https://doi.org/10.1021/acscentsci.8b00932.

(32) Tan, M. L.; Ling, L.; Fischbach, C. Engineering Strategies to Capture the Biological and Biophysical Tumor Microenvironment in Vitro. Adv. Drug Deliv. Rev. 2021, 176, 113852. https://doi.org/10.1016/j.addr.2021.113852.

(33) Qu, Y.; Han, B.; Yu, Y.; Yao, W.; Bose, S.; Karlan, B. Y.; Giuliano, A. E.; Cui, X. Evaluation of MCF10A as a Reliable Model for Normal Human Mammary Epithelial Cells. PLoS One 2015, 10 (7), 1–16. https://doi.org/10.1371/journal.pone.0131285.

(34) Samson, J.; Derlipanska, M.; Zaheed, O.; Dean, K. Molecular and Cellular Characterization of Two Patient-Derived Ductal Carcinoma in Situ (DCIS) Cell Lines, ETCC-006 and ETCC-010. BMC Cancer 2021, 21 (1), 1–20. https://doi.org/10.1186/s12885-021-08511-2.

(35) Vidavsky, N.; Kunitake, J. A.; Chiou, A. E.; Northrup, P. A.; Porri, T. J.; Ling, L.; Fischbach, C.; Estroff, L. A. Studying Biomineralization Pathways in a 3D Culture Model of Breast Cancer Microcalcifications. Biomaterials 2018, 179, 71–82. https://doi.org/10.1016/j.biomaterials.2018.06.030.

(36) Madupalli, H.; Pavan, B.; Tecklenburg, M. M. J. Carbonate Substitution in the Mineral Component of Bone: Discriminating the Structural Changes, Simultaneously Imposed by Carbonate in A and B Sites of Apatite. J. Solid State Chem. 2017, 255, 27–35. https://doi.org/10.1016/j.jssc.2017.07.025.

(37) Doherty, W. O. S.; Crees, O. L.; Senogles, E. The Preparation of Calcium Oxalate Dihydrate Crystals. Cryst. Res. Technol. 1994, 29 (4), 517–524. https://doi.org/10.1002/crat.2170290412.

(38) Yuhas, J. M.; Li, A. P.; Martinez, A. O.; Ladman, A. J. A Simplified Method for Production and Growth of Multicellular Tumor Spheroids. Cancer Res. 1977, 37 (10), 3639–3643.

(39) Schneider, C. A.; Rasband, W. S.; Eliceiri, K. W. NIH Image to ImageJ: 25 Years of Image Analysis. Nat. Methods 2012, 9 (7), 671–675. https://doi.org/10.1038/nmeth.2089.

(40) Iqbal, N.; Iqbal, N. Human Epidermal Growth Factor Receptor 2 (HER2) in Cancers: Overexpression and Therapeutic Implications. Mol. Biol. Int. 2014, 2014, 1–9. https://doi.org/10.1155/2014/852748.

(41) Loibl, S.; Gianni, L. HER2-Positive Breast Cancer. Lancet 2017, 389 (10087), 2415–2429. https://doi.org/10.1016/S0140-6736(16)32417-5.

(42) Mitri, Z.; Constantine, T.; O’Regan, R. The HER2 Receptor in Breast Cancer: Pathophysiology, Clinical Use, and New Advances in Therapy. Chemother. Res. Pract. 2012, 2012, 1–7. https://doi.org/10.1155/2012/743193.

(43) Dean-Colomb, W.; Esteva, F. J. Her2-Positive Breast Cancer: Herceptin and Beyond. Eur. J. Cancer 2008, 44 (18), 2806–2812.https://doi.org/10.1016/j.ejca.2008.09.013.

(44) Huang, M. Y.; Chaturvedi, L. S.; Koul, S.; Koul, H. K. Oxalate Stimulates IL-6 Production in HK-2 Cells, a Line of Human Renal Proximal Tubular Epithelial Cells. Kidney Int. 2005, 68 (2), 497–503. https://doi.org/10.1111/j.1523-1755.2005.00427.x.

(45) Jeong, B. C.; Kwak, C.; Kyu, S. C.; Bong, S. K.; Sung, K. H.; Kim, J. I.; Lee, C.; Hyeon, H. K. Apoptosis Induced by Oxalate in Human Renal Tubular Epithelial HK-2 Cells. Urol. Res. 2005, 33 (2), 87–92. https://doi.org/10.1007/s00240-004-0451-5.

(46) Castellaro, A. M.; Tonda, A.; Cejas, H. H.; Ferreyra, H.; Caputto, B. L.; Pucci, O. A.; Gil, G. A. Oxalate Induces Breast Cancer. BMC Cancer 2015, 15 (1), 761. https://doi.org/10.1186/s12885-015-1747-2.

(47) Lee, W. H.; Loo, C. Y.; Rohanizadeh, R. A Review of Chemical Surface Modification of Bioceramics: Effects on Protein Adsorption and Cellular Response. Colloids Surfaces B Biointerfaces 2014, 122, 823–834. https://doi.org/10.1016/j.colsurfb.2014.07.029.

(48) Jeong, J.; Kim, J. H.; Shim, J. H.; Hwang, N. S.; Heo, C. Y. Bioactive Calcium Phosphate Materials and Applications in Bone Regeneration. Biomater. Res. 2019, 23 (1), 4. https://doi.org/10.1186/s40824-018-0149-3.

(49) Rimer, J. D.; Wesson, J. A.; Ward, M. D. Pathological Biomineralization of Calcium Oxalate Kidney Stones. In AIChE Annual Meeting, Conference Proceedings; 2008; Vol. 3, pp 415–421. https://doi.org/10.2113/GSELEMENTS.3.6.415 K.

(50) Kanlaya, R.; Sintiprungrat, K.; Thongboonkerd, V. Secreted Products of Macrophages Exposed to Calcium Oxalate Crystals Induce Epithelial Mesenchymal Transition of Renal Tubular Cells via RhoA-Dependent TGF-B1 Pathway. Cell Biochem. Biophys. 2013, 67 (3), 1207–1215. https://doi.org/10.1007/s12013-013-9639-z.

(51) Sun, X. Y.; Xu, M.; Ouyang, J. M. Effect of Crystal Shape and Aggregation of Calcium Oxalate Monohydrate on Cellular Toxicity in Renal Epithelial Cells. ACS Omega 2017, 2 (9), 6039–6052. https://doi.org/10.1021/acsomega.7b00510.

(52) Sun, X. Y.; Ouyang, J. M.; Yu, K. Shape-Dependent Cellular Toxicity on Renal Epithelial Cells and Stone Risk of Calcium Oxalate Dihydrate Crystals. Sci. Rep. 2017, 7 (1), 1–13. https://doi.org/10.1038/s41598-017-07598-7.

