## Supplementary material for "Microcalcifications can either trigger or suppress breast precancer malignancy potential according to the mineral type in a 3D tumor model": SI

Amit Cohen[1], Lotem Gotnayer[1], Dina Aranovich[1], Netta Vidavsky[1,2]
[1] Department of Chemical Engineering, Ben-Gurion University of the Negev, Beer Sheva
[2] Ilse Katz Institute for Nanoscale Science & Technology, Ben-Gurion University of the Negev, Beer Sheva

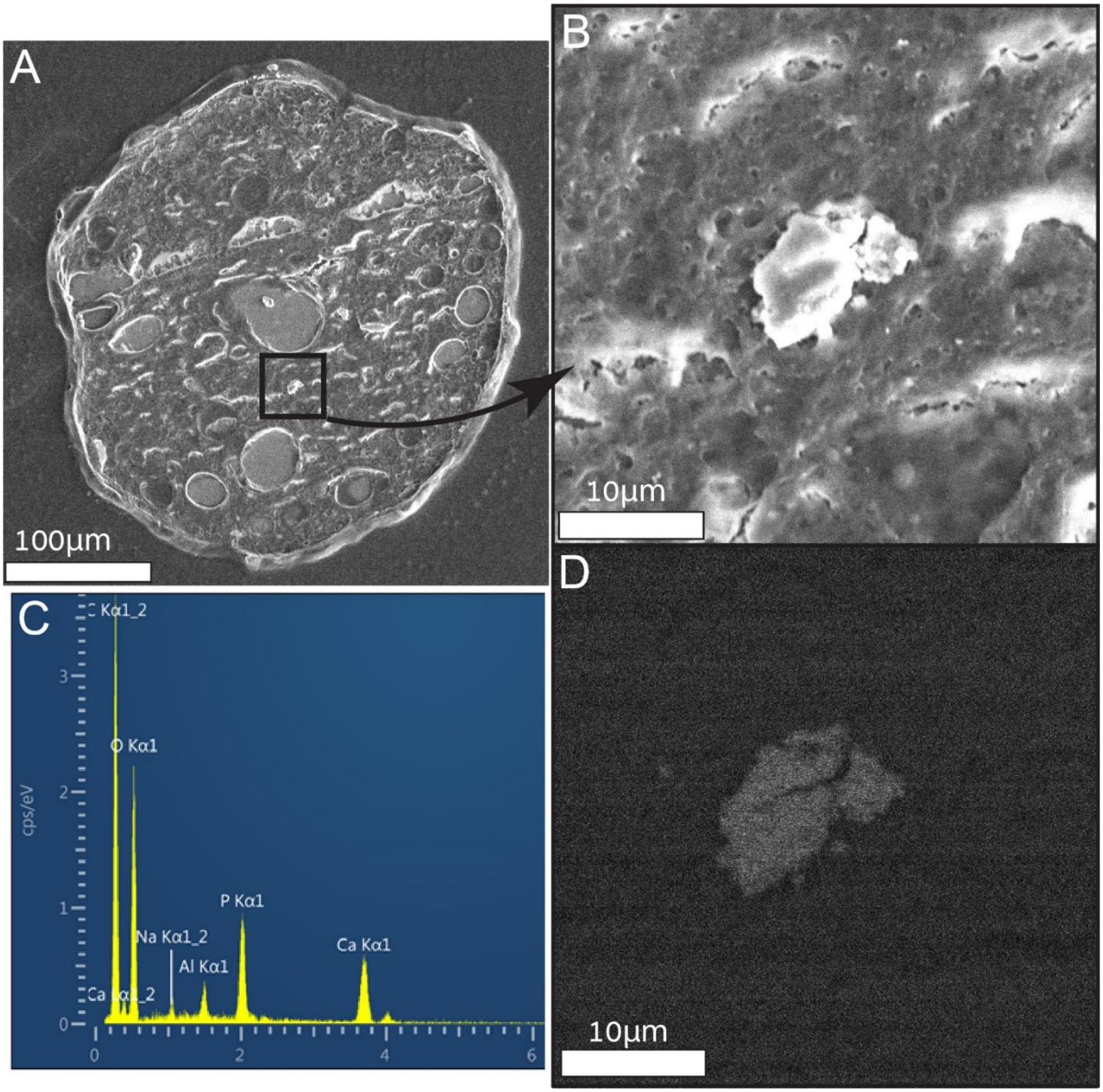


**Figure S1. A**. An SEM image of a DCIS spheroid section showing embedded MC analogs. **B.** Magnification of an MC analog in the spheroid section. **C.** EDS analysis showing Ca and P atoms, indicating that the MC consists of calcium phosphate. **D.** Back-scattered electron image of the MC demonstrating how the mineral particles, which appear as bright particles compared to their surrounding organic material, are located during SEM imaging.

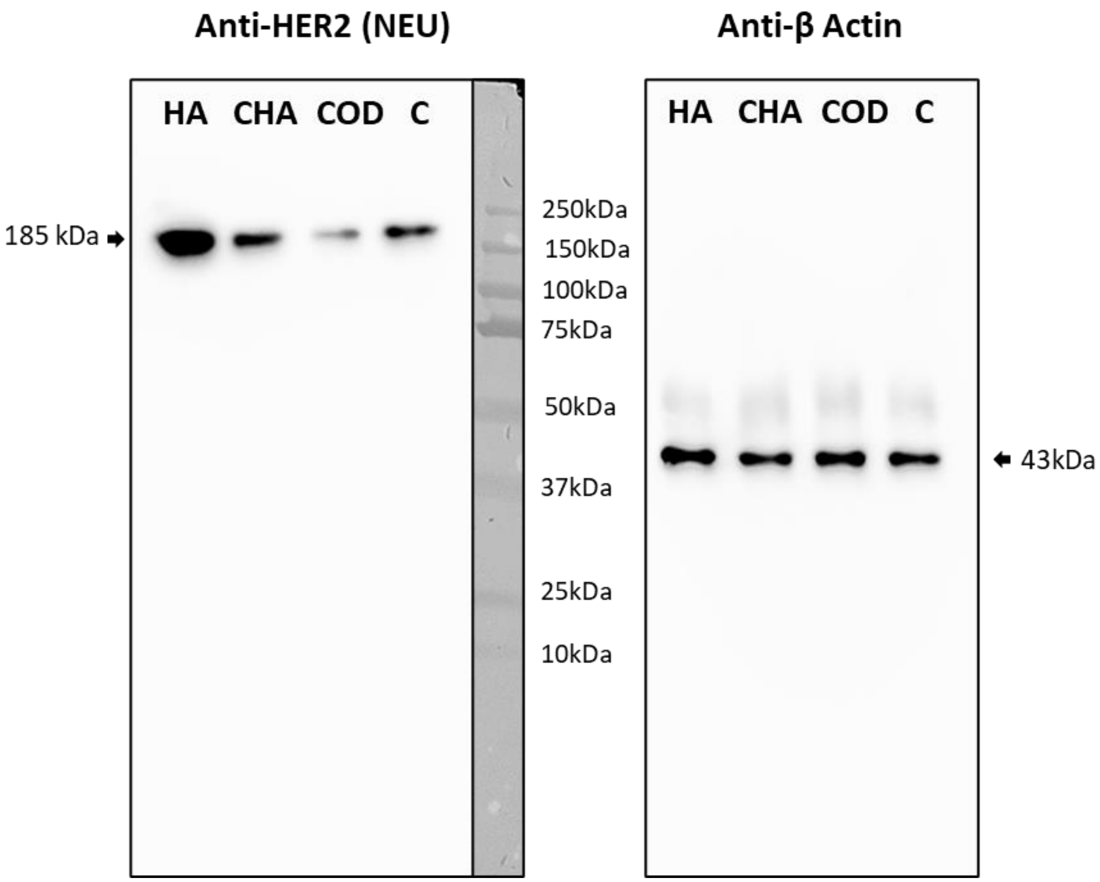


**Figure S2**. Western blot with anti Her2 and anti β-Actin antibodies. The four samples were loaded in two duplicates on the same gel. After the transfer, the membrane was cut into two halves. One half was incubated with anti – HER2 (NEU) diluted at 1:50, and the second half was incubated with anti – β-Actin diluted (1:400). HA=low carbonate apatite and CHA=high carbonate apatite.
